# Sim3C: simulation of HiC and Meta3C proximity ligation sequencing technologies

**DOI:** 10.1101/134452

**Authors:** Matthew Z DeMaere, Aaron E Darling

## Abstract

**Background:** Chromosome conformation capture (3C) and HiC DNA sequencing methods have rapidly advanced our understanding of the spatial organization of genomes and metagenomes. Many variants of these protocols have been developed, each with their own strengths. Currently there is no systematic means for simulating sequence data from this family of sequencing protocols.

**Findings:** We describe a computational simulator that, given reference genome sequences and some basic parameters, will simulate HiC sequencing on those sequences. The simulator models the basic spatial structure in genomes that is commonly observed in HiC and 3C datasets, including the distance-decay relationship in proximity ligation, differences in the frequency of interaction within and across chromosomes, and the structure imposed by cells. A means to model the 3D structure of topologically associating domains (TADs) is provided. The simulator also models several sources of error common to 3C and HiC library preparation and sequencing methods, including spurious proximity ligation events and sequencing error.

**Conclusions:** We have introduced the first comprehensive simulator for 3C and HiC sequencing protocols. We expect the simulator to have use in testing of HiC data analysis algorithms, as well as more general value for experimental design, where questions such as the required depth of sequencing, enzyme choice, and other decisions must be made in advance in order to ensure adequate statistical power to test the relevant hypotheses.

## Findings

### Software testing

Within bioinformatics, data simulation has become an important proxy for real experimental data when testing individual algorithms and, more broadly, whole analysis workflows. Use of simulated data in testing can be motivated by the formal notion of software validation, the direct comparative analysis of algorithms or even as an exploratory technique when prototyping experimental design. For established experimental methods, the ever-accumulating public data archive offers a route to thorough real-data driven testing. For a chosen test however, a difficulty remains in matching desired data characteristics to one or several public dataset(s). Further, as fields such as DNA sequencing develop, new forms of experimental data appear for which the public data archives contain few, if any, examples. Though performance on real data is the ultimate arbiter of analytical value, few researchers would have the time and financial resources to commit to its generation purely for software testing. Simulation-driven development and testing has proven to be a highly cost effective and time efficient approach. It offers the possibility to explore data characteristics as a near continuum and subject software to a previously unavailable degree of testing thoroughness.

Tools for simulating DNA sequencing reads have existed from the very early days of genomics, beginning with the many anonymous implementations of simple DNA shearing algorithms, up to the most recent highly detailed empirical model simulators [13, 14, 19, 30]. From read simulation in isolation, field advancements such as metagenomics have been accompanied soon after by simulators reflecting their specific data characteristics and evolving experimental methodology [2, 16, 34].

We introduce Sim3C, a software package designed to simulate data generated by HiC and other 3C-based proximity ligation (PL) sequencing protocols. The software includes flexible support for a range of sequencing project scenarios and choice of three 3C methods (HiC, Meta3C, DNase-HiC). The resulting output (paired-end FastQ) is easily assimilated into existing workflows, opening the door to more thorough software testing, such as the comparative analysis of clustering algorithms [8].

### 3C sequencing

3C-based sequencing protocols, including Hi-C, 4C-seq, and Meta3C, have great potential to address questions directed at the spatial organization of DNA in samples ranging from eukaryotic tissue, to single cells, to microbial communities. The growing use of these protocols creates a legitimate need for a simulator capable of generating data with relevant characteristics.

Chromosome conformation capture (3C) was originally designed as a PCR-based assay to measure interactions among a small number of defined regions of eukaryotic chromosomes [7]. In 2009 Lieberman-Aiden [21] reported an extension of the protocol to high throughput sequencing, enabling the global spatial arrangement of chromosomes to be reconstructed at unprecedented resolution. All 3C protocols depend on an initial formalin fixation step, which crosslinks proteins bound to DNA in vivo. Subsequently cells are lysed and the DNA:protein complexes are sheared enzymatically and/or physically to create free ends in the bound DNA strands. These free ends are then subjected to a proximity ligation reaction, in which ligation of free ends preferentially occurs among DNA strands cobound in a protein complex. The DNA:protein crosslinks are then reversed, the DNA is purified, and an Illumina-compatible sequencing library is constructed. In HiC protocols, the proximity ligation junctions can then be further purified in the sequencing library.

3C-derived methods have found several applications beyond their initial use to reconstruct 3D chromosome structure. For example, it has been shown that 3C-derived data provide a valuable signal for genome scaffolding [5, 10], as well as a signal that can support genome-wide haplotype phasing [17, 36]. 3C-derived data has also proven valuable for metagenomics, where initial studies on mock communities demonstrated that highly accurate genome reconstruction in mixed microbial communities could be facilitated by proximity ligation sequence data [4, 6, 26]. Subsequent application to naturally occurring microbial communities has also suggested that bacteriophage can be linked to their hosts with this data type [24].

In the remainder of this manuscript we describe the Sim3C software and outline how it can be used to simulate data for various 3C-derived experiments.

### Experiment scenarios

Beyond simple monochromosomal genome sequencing experiments, Sim3C offers support for the more complex scenarios of multi-chromosomal genomes and metagenomes. A scenario is defined by way of a community profile; assigning a copy-number and containing genome to each chromosome and a relative abundance to each genome. The profile and supporting reference sequences form a skeleton definition with which to initialize the weighted random sampling process within a simulation. The user can elect to supply a profile either as an explicit table (listing 1, 2) or allow Sim3C to draw abundances at runtime from one of three distributions (equal abundance, uniformly random, log-normal distribution) for communities made up of strictly mono-chromosomal genomes.

**Listing 1.**
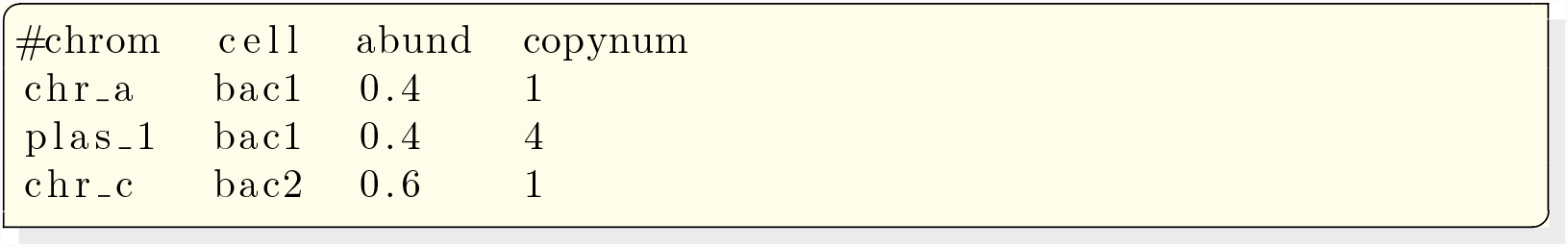
A mock two genome community. For demonstration purposes, we assume that the plasmid (plas_1) is present in four copies and that there is a 0.4/0.6 relative abundance split between the two organisms (bac1, bac2) in the community

**Listing 2.**
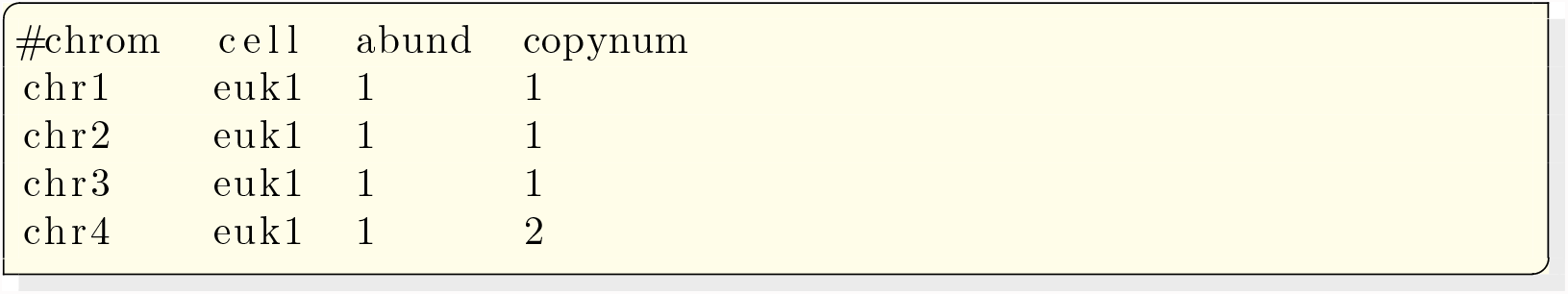
A mock four chromosome genome. Cellular abundance is a constant across the profile, while chr4 exists in two copies. Note that relative abundances specified in a profile are not required to sum to 1, but are normalised internally.

### Error Modelling

Sim3C models three forms of experimental noise: machine-based sequencing error, the formation of spurious ligation products and the contamination of PL libraries with WGS read-pairs.

To simulate machine-based sequencing error, the paired-end mode from art illumina [14] has been reimplemented as a Python module (`Art.py`). This approach was taken as delegating read-pair generation to native invocations of art_illumina proved cumbersome. More explicitly, a loosely coupled solution (via subprocess calls but without an IPC mechanism) lacked sufficient control to generate PL read-pairs in an efficient and robust manner. On the other hand, tightly coupling Sim3C to the ART C/C++ source code (i.e. implementing hooks) would have left Sim3C vulnerable to changes in a non-public external API (i.e. a codebase without formal definition or guarantee of stability). Reimplementation also meant Art's many empirically derived machine profiles are available for use by Sim3C, allowing equivalent treatment of machine-error when experiments involve both PL (Sim3C) and pure WGS (art_illumina) libraries.

The production of spurious ligation products is an inherent source of noise in PL library construction [28]. Sim3C models spurious pairs as the uniformly random ligation of any two cut-sites across all source genomes. While this process disregards cellular organisation, it respects the relative abundance of chromosomes. Spurious pairs, and to a lesser extent sequencing error, represent an important confounding signal to downstream analyses that attempt to infer the cellular or chromosomal organisation of DNA sequences.

Lastly, conventional WGS read-pairs represent a source of contamination within a PL library, which even after HiC enrichment steps, are not completely eliminated. The rates at which spurious and WGS read-pairs are injected into a simulation run are controllable by the end-user.

### Simulation modes

Since HiC was first introduced [21], the development of variants and extensions has been continual [11, 26, 32, 33]. Variants have often strived to further enhance the discriminatory power of the original experiment, while seemingly adding yet more complexity to an already challenging protocol (*in-situ* DNase HiC, sciHiC) [33]. Others instead have sought compromise, with the aim of lessening the burden on the laboratory (Meta3C). While not considering more recent and complex extensions, Sim3C offers three simulation modes: traditional HiC, Meta3C and DNase-HiC. The first two of these modes were chosen as representing the fundamental basis (traditional HiC) and an attractive and pragmatic simplification of the original (Meta3C). The third mode (DNase-HiC) replaces the restriction endonuclease driven production of the free-ends, used to form PL products, with an ideally-free process of DNA fragmentation. In the laboratory, this ideally-free process could be carried out by DNase digestion or mechanical shearing via sonication.

The most notable difference between the methods of HiC and the more recent Meta3C, is that after restriction digest, HiC employs additional steps leading to the incorporation of biotin tags at each PL junction. This biotinylation permits HiC libraries to be subsequently enriched for fragments containing PL junctions by streptavidin-mediated affinity purification. Without enrichment, the simpler Meta3C protocol results in a gross mixture of both WGS and PL read-pairs, where only a small percentage of the total read-pair yield (approx. 1%) will possess PL junctions [22]. The enrichment process within HiC, however, is not perfectly efficient and WGS read-pairs are still observed (approx. 10–50% of reads contain a PL product) [22]. DNase-HiC replaces restriction digest with a non-specific endonuclease (e.g. DNase I) [23] or mechanical DNA shearing process (e.g. sonication) [11]. In this operational mode, Sim3C treats DNA cleavage as a completely unbiased (free) process and as such all genomic positions have equal probability of participating in proximity ligation events.

Within Sim3C, each of the three methodological variations is conceptualised as a sequencing strategy (figure 1) and each iteration of a strategy produces one read-pair (PL or WGS in origin). For all strategies, an iteration begins by drawing a 3-tuple of insert parameters: length, direction and junction point (*L_ins_, dir, x_junc_)*.

**Figure 1.**
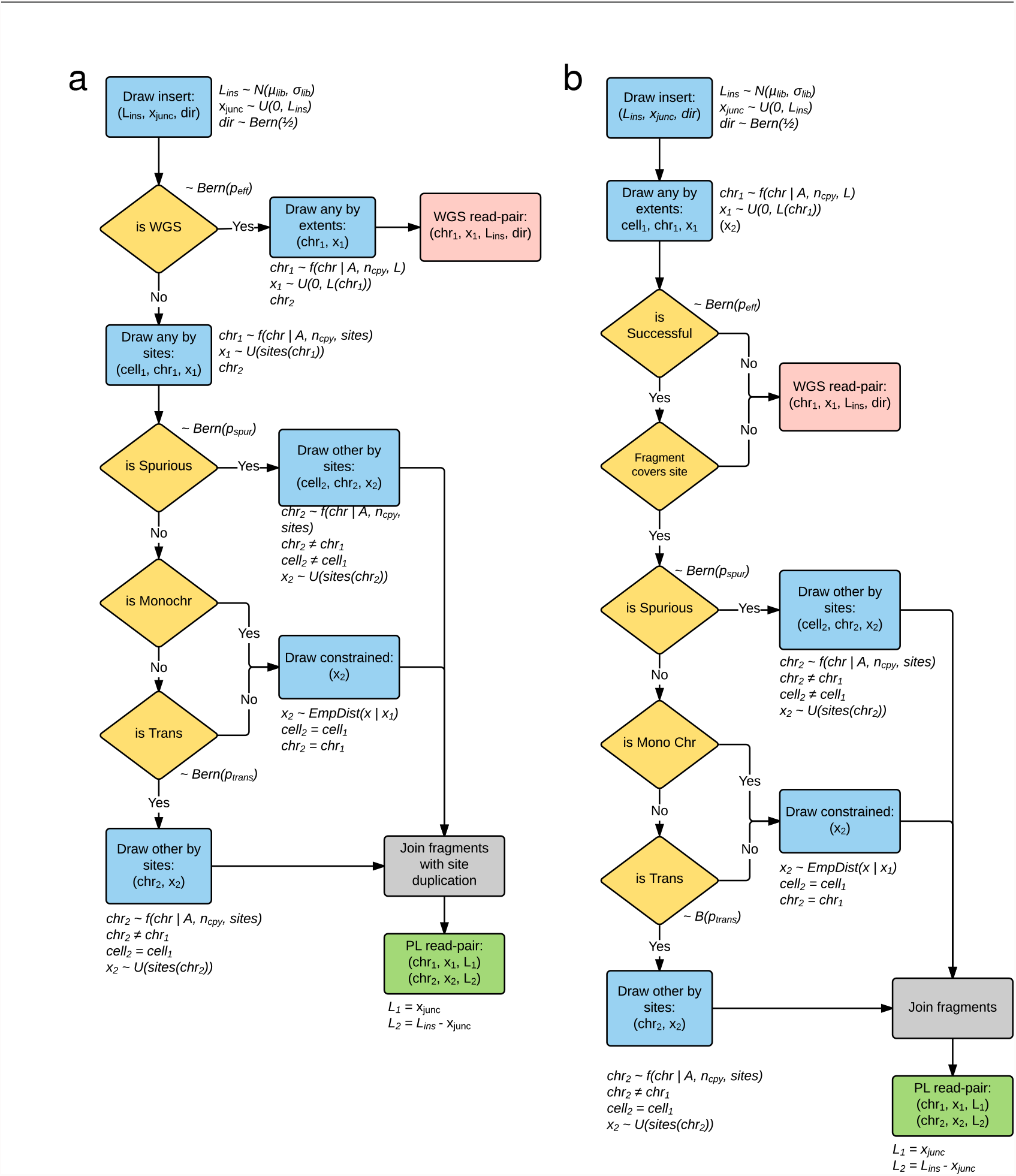
Logical schema used within Sim3C. (**a**) HiC and (**b**) Meta3C simulat strategies. Gold diamonds represent simple Bernoulli trials. Blue boxes repres sampling distributions defined by runtime input data (community profile, geno sequences, enzyme) and the empirically derived distribution for intra-chromosome (*cis*) interaction probability (equation 1). Logical end-points to a single iteration of eit algorithm are represented as red (producing a WGS read-pair) and green boxes (produc a PL read-pair). Due to the elimination of the biotinylation step, Meta3C does produce a duplication of the restriction cut-site overhang (grey boxes).

After obtaining insert parameters, the HiC strategy (figure 1a) first tests if the insert will represent a WGS or PL read-pair (~*Bern*(*p_eff_*)), where efficiency *p_eff_* is defined in the sense of enrichment. When *p_eff_* = 1, there is perfect filtering and all WGS read-pairs are eliminated from the experiment. In the case of WGS, the iteration reaches an end-point and the simulation emits a conventional read-pair drawn from the community definition. In the case of PL, next a 3-tuple defining a cut-site is drawn (*gen*_1_, *chr*_1_, *x*_1_), where the categorical distribution over chromosomes is weighted by relative abundances (*A*) and chromosomal copy-numbers (*n_cpy_*); genomic position is sampled uniformly from the set of restriction sites (*sites*(*chr*_1_)); and parent genome (*gen*_1_) is implicit from the chromosome. Next, a test for spurious ligation is performed (~*Bern*(*p_spur_*)). If a spurious event has occurred, the 3-tuple defining the second site (*gen*_2_, *chr*_2_, *x*_2_) is drawn i.i.d. as the first. If not spurious, next a test for inter-chromosomal (*trans*) ligation is performed. For inter-chromosomal events, as the source genome is implicitly defined (*gen*_2_ = *gen*_1_), only the second chromosome and position (*chr*_2_, *x*_2_) are drawn. Chromosome chr2 is selected without replacement from the same genome (*gen*_1_), where the categorical distribution is adjusted for the removal of the first chromosome, and genomic position *x*_2_ on *chr*_2_ is drawn i.i.d. as the first. Lastly, an intra-chromosomal (*cis*) ligation must have occurred. As now both genome and chromosome are implicitly defined (*gen*_2_ = *gen*_1_, *chr*_2_ = *chr*_1_), all that is required is to draw genomic position *x*_2_. Here, the separation *s* between *x*_1_ and *x*_2_ (*s* = |*x*_2_ − *x*_1_|) is constrained to follow an empirically determined long-tailed mixture of the geometric and uniform distributions (equation 1).

For Meta3C (figure 1b) after insert parameters are determined, in the same fashion as a regular WGS read, an initial free genomic position is drawn 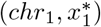, uniformly distributed over the extent of *chr*_1_ rather than only over its cut-sites. In real datasets, it has been observed that neither the restriction digestion nor the re-ligation of free ends are perfectly efficient. Taken as independent probabilities, in our model we conceptualise their joint occurrence as an efficiency factor, *p_eff_* and a Bernoulli trial (*Bern*(*p_eff_*)) determines whether a sequence read is successful in containing an observable proximity ligation event. Failing this coverage test relegates the iteration and end-point and emit a WGS read-pair. Successful candidates instead continue akin to the HiC decision tree, beginning with the test for spurious ligation.

For both HiC and Meta3C, PL read-pairs are produced by joining the free-ends drawn above as defined by the fragment parameters (figure 2). Here the location of the PL junction within the insert is determined by *x_junc_*. At the junction, HiC differs from Meta3C as the process of biotinylation results in the duplication of the restriction cut-site overhang sequence. The overhang duplication in Hi-C is included in the simulation.

**Figure 2.**
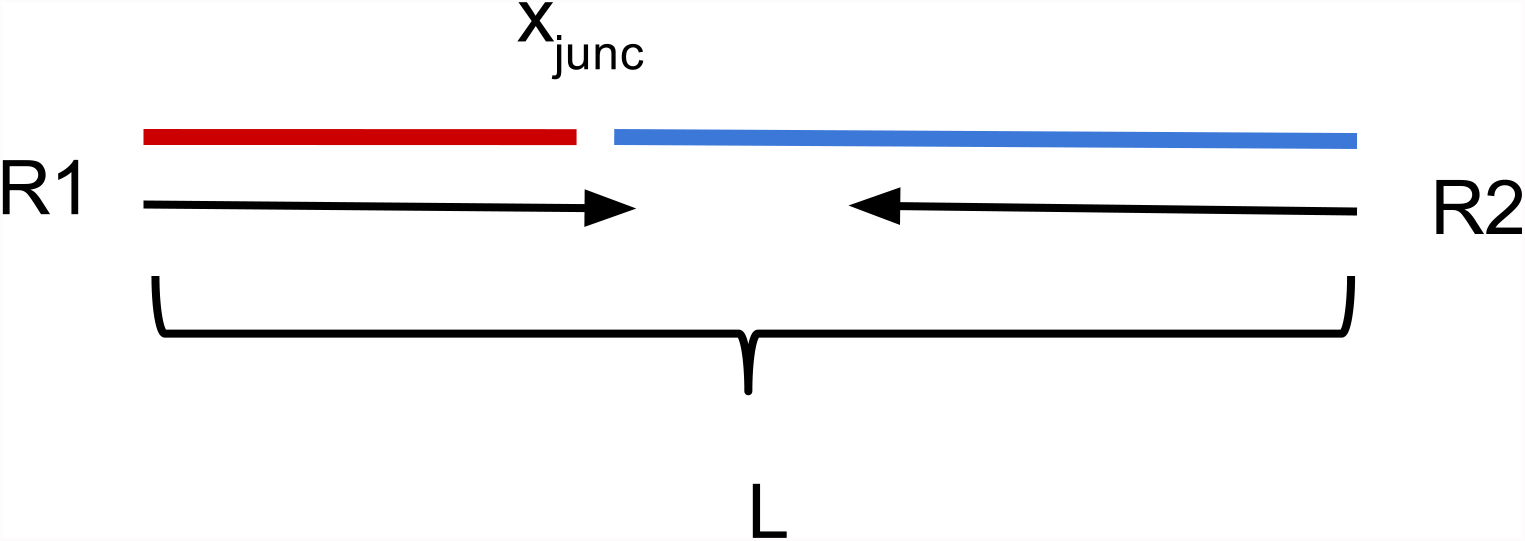
Proximity ligation insert. The joining of parts A (red) and B (blue) to create the full junction containing PL insert, from which the read-pair is then simulated via Art.py. Here, the junction point varies uniformly over the interval [0*..L*) and in combination with the requested read length and insert size range, can result in read-through events as are observed in real experiments. In the case of traditional HiC, the side-effect of site duplication makes it possible to confirm such occurrences.

DNase-HiC is handled similarly to traditional HiC, with the exception that, as *in-silico* digestion trivially leads to all sites, the simulated digestion is unnecessary to perform and positions can be drawn directly from the uniform distribution over the interval [0..*L_chr_*). Site duplication, attributable to the likely production of random overhangs in this scenario, is not presently simulated.

### Empirical distribution

The central relation used to express the probability of observing intra-chromosomal proximity ligation (PL) events as a function of genomic separation *s* is a mixture model combining the geometric and uniform distributions (equation 1), with mixing parameter *β*, geometric distribution shape parameter *α* and genomic interval length *l*. This combination allows for a non-zero probability for all chromosomal positions, while also capturing the rapid early fall-off in probability with increasing separation (figure 3). Agreement with empirical observations is acceptable over the majority of the range, diverging slightly from the empirical estimate at small separation (*s* < 10 kbp). Worth considering, however, is that as diminishing separation *s* approaches the length of the sequencing insert, there is increasing odds of PL read-pair counting statistics being contaminated by WGS read-pairs that escaping filtering efforts.

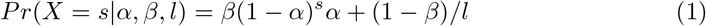

**Figure 3.**
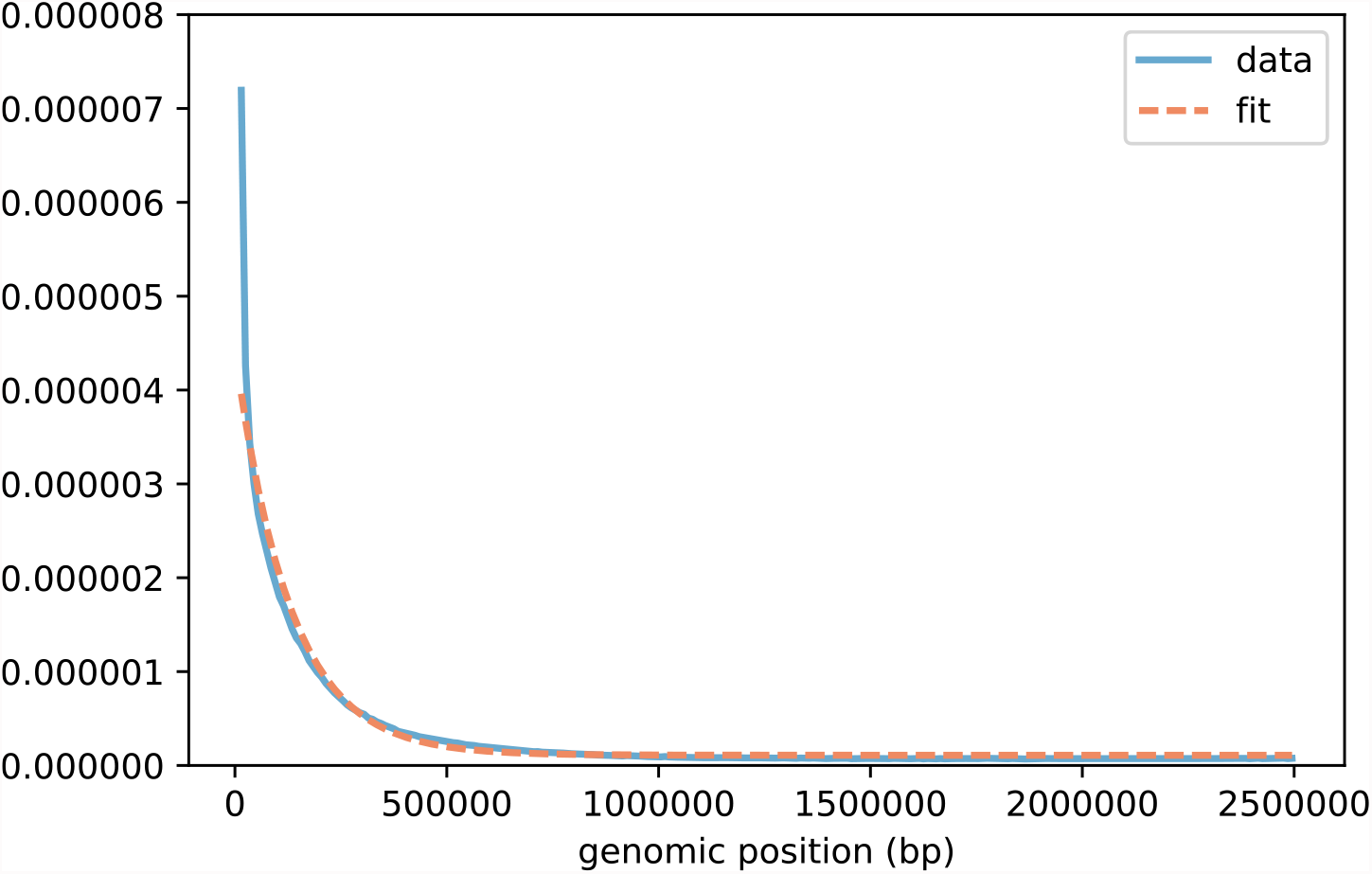
Empirical model against real data. Agreement of model relation (equation 1) as fitted to empirical data collected from the single bacterial genome *C*. *crescentus* (*α*=7.7e-6, *β*=0.56). Only the first 2.5Mbp are shown so as to see the highest density region near for small separation.

**Figure 4.**
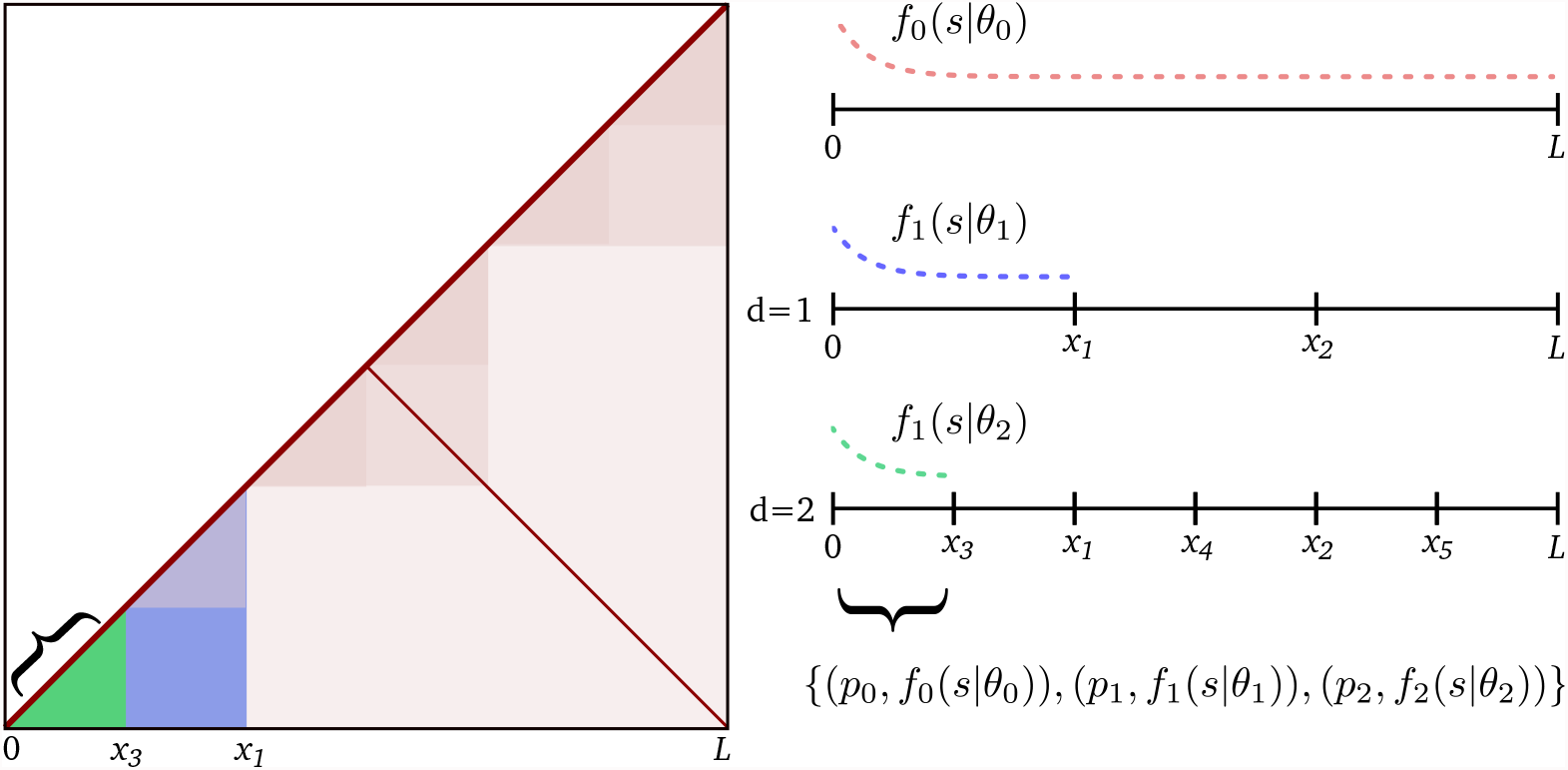
Approximating the structurally related modulation of observed contact frequencies. Beyond primary intra-chromosomal interactions which produce the main diagonal, Sim3C can reproduce an inter-arm mediated anti-diagonal at user controlled intensity. Illustrated here, finer scale modulations attributed to topologically associated domains (TADs) can optionally be approximated. Primary interactions are governed by the empirical distribution f0(s—0) which covers the full interval [0*, L*). Each level of recursion (*d* = 1, 2 ⋯ *n*) generates a finer set of intervals, to which a distribution *f_i_* (*s*|*θ_i_*) and probability *p_i_* is assigned. The intervals at the final depth each define a range (green, curly braces) over which a set of probabilities and empirical distribution pairs govern interaction separation *s*.

### Structurally related interactions

Sim3C can approximate the modulation of observed contact frequencies brought about by 3D chromosome structure. Independent of any 3D structure that might exist, the primary and most frequently observed interactions are those which occur along a chromosome (intra-arm) (figure 5a), seen as the primary (*y* ≃ *x*) diagonal in the con map. Less intense are interactions occurring between chromosomal arms (inter-arm (figure 5b) [18], which produced an anti-diagonal (*y* ≃ *L* − *x*) in the contact map. At progressively smaller scales, the hierarchical 3D folding of DNA into topologically associated domains (TADs) produces overlapping regions of interaction (figure 5c) visible in the contact map as block-like intensity modulations. Though the agents responsible for their formation vary [1, 3], the characteristic patterns evident in real-data derived 3C contact maps have been observed across all three domains [9, 18, 37].

**Figure 5.**
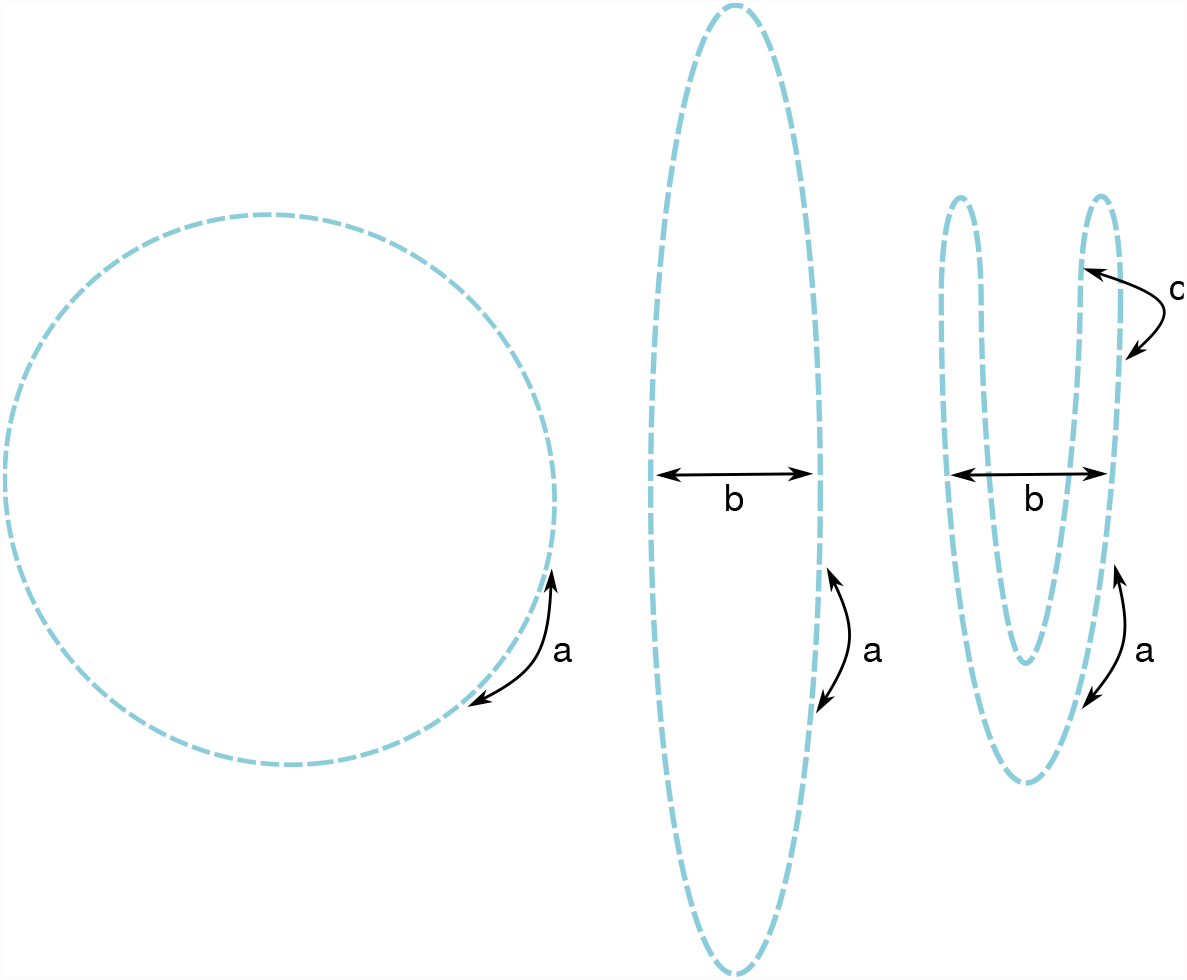
Interactions from simple folds. For a flexible and unbound chromosome (circular here) embedded in 3D space, the possibility of spatially proximate interactions can be increased from (**a**) those which are strictly primary, to (**b**) include inter-arm and (**c**) further still by successive folding. When the spatial arrangement is consistent across the population of cells, this will be observable as modulations in the contact frequencies.

Our approximation of hierarchical folding begins from the full extent *L* of a chromosome (figure 4). Folding is portrayed by the division of the interval [0..*L*) in set of non-overlapping sub-intervals {[0, *x*_1_), [*x*_1_, *x*_2_),⋯, [*x*_*n*-1_, *x_n_*)}, the number a widths of which are drawn at random (*U*(*l_min_*, *l_max_*), *U*(*n_min_*, *n_max_*)). The procedur then recursively applied to each sub-interval until a depth *d*, producing a nested se coverings of the full interval [O..*L*) at progressively finer scales. Across this hierarch collection each interval is assigned a uniformly distributed random probability *P_i_* a empirical distribution *f_i_*(*s*|*θ_i_*) (equation 1) for separations parameterised by shape parameter *α_TAD_* and interval length *l_inv_* = *x*_*i*+1_ − *x_i_*, where *θ* = (*α_TAD_, β, l_inv_*).

The process of drawing samples of separation begins by determining the set of intervals {*l_inv_*} which contain an initial point *x*_0_. The intervals, as tuples (*p_i_*, *f_i_* (*s*|*θ_i_*)), then form a categorical distribution (equation (7)), from which a governing distribution *f_i_* (*s*|*θ_i_*) is drawn and finally a sample of separation is taken, *s* ~ *f_i_* (*s*|*θ_i_*). To efficiently sample from the full collection, an interval-tree data structure is employed. When queried, an interval-tree returns the set of intervals {l} overlapping a position *x* in order *O*(log *n* + *m*), where *n* is number of intervals and *m* is number of intervals returned by the query.

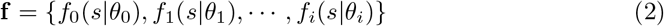

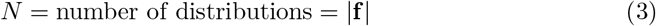

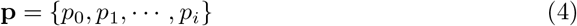

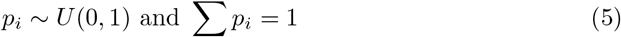

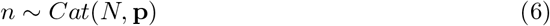

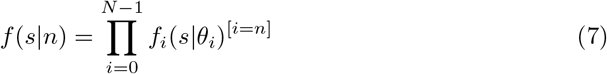

where [*i* = *n*] is the Iverson bracket.

### Example scenarios

In the following, three use-cases are presented to demonstrate aspects of the resulting simulation output: bacterial genome, multi-chromosomal eukaryotic (yeast) genome, and metagenome. For each use-case, 3C contact maps have been used to pit simulation output against the corresponding real experimental data (table 1).

**Table 1.**
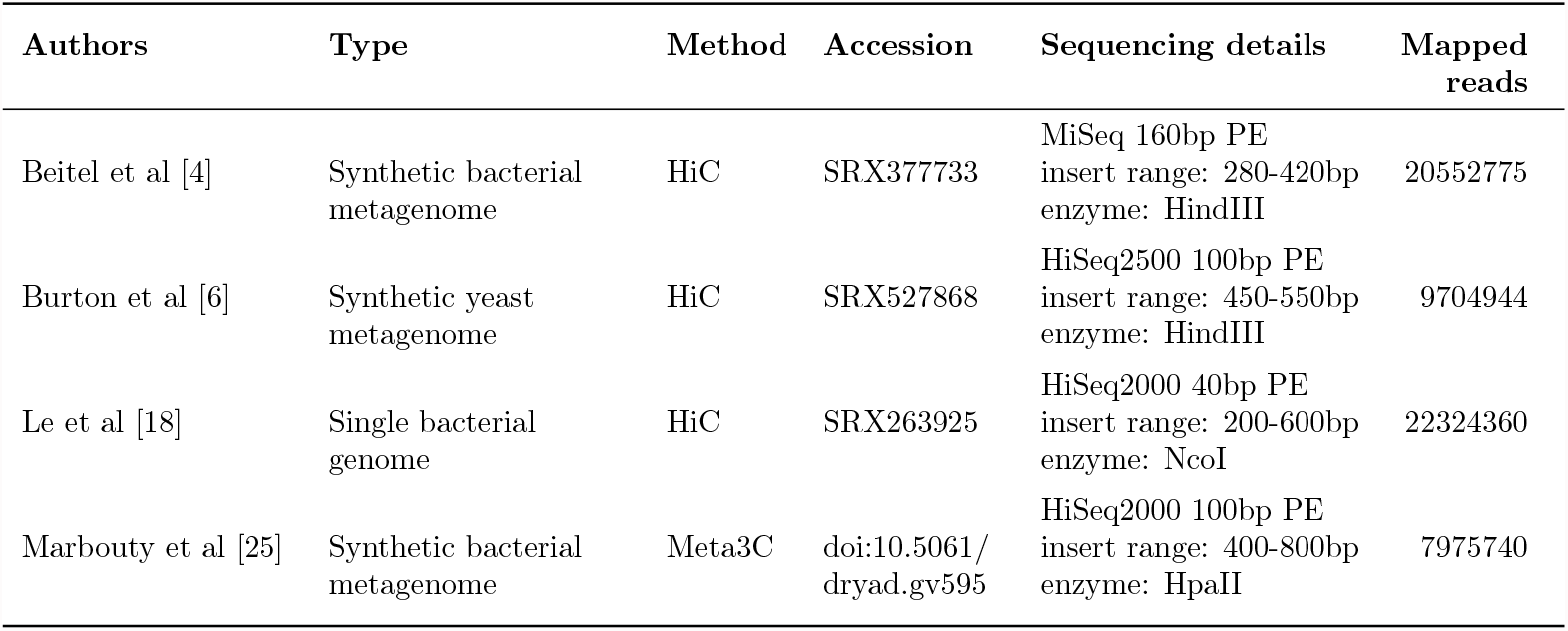
Real HiC and Meta3C data-sets used within this work. The total off-diagonal weight of the contact map was used to calibrate the amount of simulated sequencing required to approximately match the outcome of the real experiments.

| Authors | Type | Method | Accession | Sequencing details | Mapped reads |
| --- | --- | --- | --- | --- | --- |
| Beitel et al [4] | Synthetic bacterial metagenome | HiC | SRX377733 | MiSeq 160bp PE<br>insert range: 280-420bp<br>enzyme: HindIII | 20552775 |
| Burton et al [6] | Synthetic yeast metagenome | HiC | SRX527868 | HiSeq2500 100bp PE<br>insert range: 450-550bp<br>enzyme: HindIII | 9704944 |
| Le et al [18] | Single bacterial genome | HiC | SRX263925 | HiSeq2000 40bp PE<br>insert range: 200-600bp<br>enzyme: NcoI | 22324360 |
| Marbouty et al [25] | Synthetic bacterial metagenome | Meta3C | doi:10.5061/dryad.gv595 | HiSeq2000 100bp PE<br>insert range: 400-800bp<br>enzyme: HpaII | 7975740 |

### Bacterial

A monochromosomal bacterial genome is perhaps the simplest scenario to which proximity ligation methods have been applied, making for a sensible entry point from which to make comparison. Due to the smaller extent, a bright and high resolution contact map (10 kbp bin size) is possible for a practical volume of sequencing data, potentially revealing fine detail not easily discerned with larger bin sizes (50-100 kbp bin size).

The genome of *Caulobacter crescentus* NA1000, a model organism in the study of cellular differentiation and regulation of the cell cycle, is comprised of a single 4 Mbp circular chromosome [27]. Deep HiC sequencing of *C. crescentus* has been used to explore the degree to which bacterial chromosomes can be regarded as organised and provided evidence for the existence of so called chromosomal interaction domains (CIDs) [18]. As a prokaryotic analog of topologically associated domains (TADs) from eukaryotic literature [1, 29, 31], these regions are believed to promote intra-domain loci interactions and thereby act to functionally compartmentalize the genome. This chromosomal structure was observed to be at once disruptable through rifampicin mediated inhibition of transcription and malleable by the movement of highly expressed genes [18].

For the raw contact map of *C. crescentus*, prominent rectilinear features are apparent for both real and simulated traditional HiC sequencing data (figure 6a,b), while notably for simulated unrestricted HiC the field is much smoother (figure 6c). Within the Sim3C model, a single distribution governs both intra- and inter-arm interactions. Inspection of the real-data contact map (figure 6a) suggests that the true relationship governing inter-arm interactions is more dispersed. This perhaps is not surprising, where different arms associating spatially possess a greater number of potential configurations than can be taken on by the primary chromosome backbone. Additionally for the real contact map, long-range interactions away from either diagonal can be seen to drop to a lower threshold than that produced from simulation.

**Figure 6.**
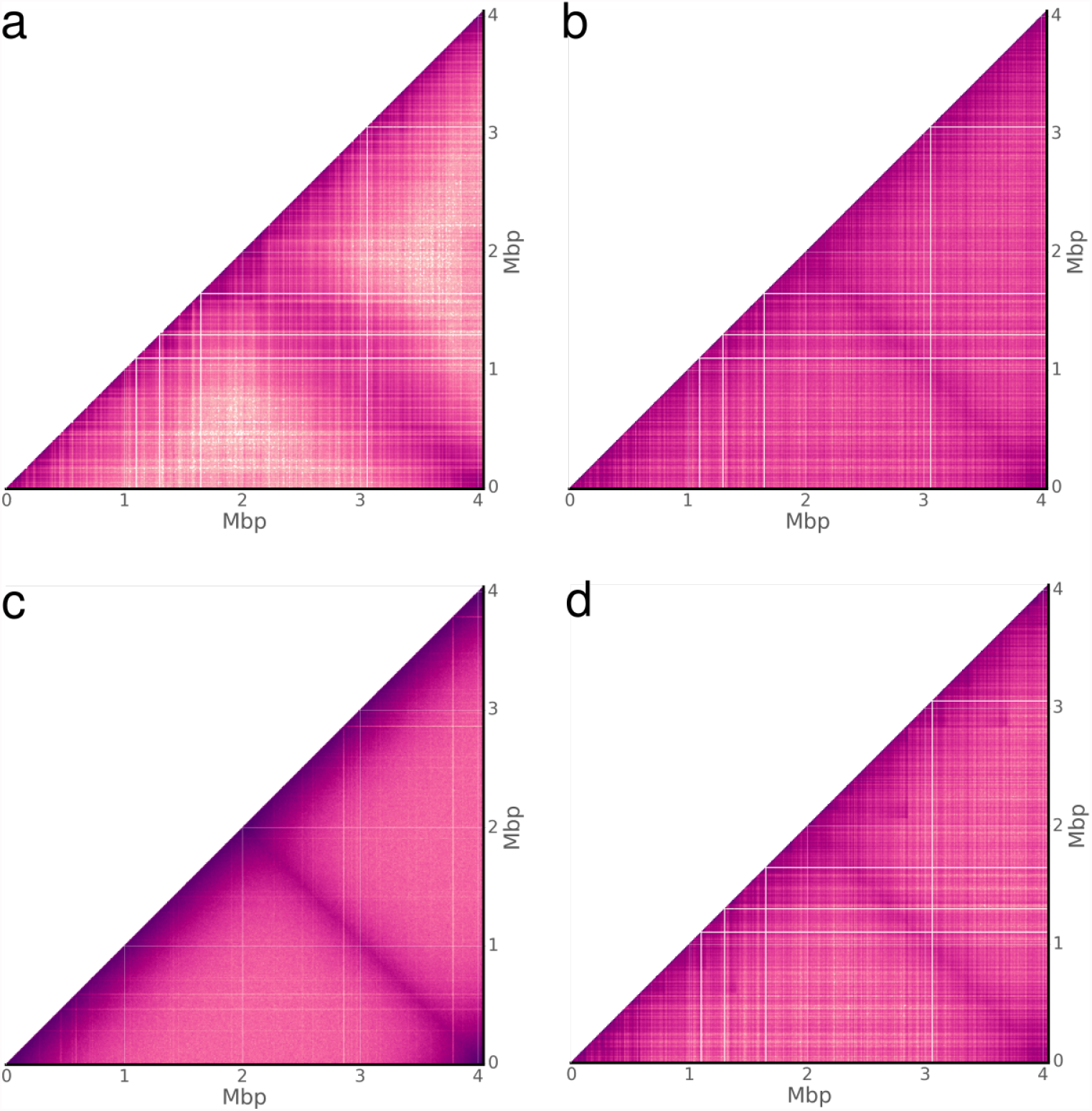
Bacterial contact maps. Observed HiC interactions for the monochromosomal genome of *Caulobacter crescentus* NA1000. Comparing (**a**) real experimental data [18], to the three simulation choices (**b**) traditional HiC, (**c**) DNase-HiC and (**d**) traditional HiC with TADs enabled. Sharp rectilinear modulations of the intensity within (**a**) and (**b**) indicate a reduction in PL observations within a given bin. Not due to 3D chromosome structure, rather such features can be attributed largely to mappability and low cut-site density. (**c**) Without an enzymatic constraint a significantly smoother field is apparent, yet still susceptible to mappability. (**d**) Enabling topologically associated domains (TADs) highlights the similarity between features produced merely from biases and what could be truly associated with 3D structure.

Within the unrestricted HiC map, the fine zero-intensity rectilinear features are a direct result of poor mappability (non-unique sequence), where their small size reflects the extent of the non-unique regions (example: rRNA genes) and the single base-pair resolution of the less constrained read generation process. The process of enzymatic digestion is the only difference between the unrestricted and traditional HiC simulation models. The clear contrast in their contact maps is thus a combination of factors either directly inherent to digestion (cut-site density) or a byproduct of downstream bioinformatics analysis (e.g. filtering heuristics). Though the problem of mappability exists for any reference based representation, for real and simulated traditional HiC, zero-intensity rectilinear features mark regions devoid of cut-sites over at least 10 kbp.

Enabling TAD approximation in simulated traditional HiC (figure 6d) has the effect of modulating map intensity in a manner not particularly distinct from that produced purely from experimental/workflow bias. Discriminating between these two feature sources; one representing experimental signal, the other representing noise; demands attention when developing solutions to problems such as normalisation. Contact map normalisation methods, whether based upon explicit or implicit bias models [35], may leave behind remnants of noise-related features from either a lack of convergence or model limitations. Downstream inferencing should therefore not be made under an assumption of bias-free signal.

### Eukaryotic

The eight chromosomes of the 15.4 Mbp genome of the native xylose-fermenting yeast *Scheffersomyces stipitis* CBS 6054 [15] range in size from 970 kbp to 3.5 Mbp. The organism was one of 16 yeasts included in a synthetic community to explore the application of HiC sequencing to deconvolving metagenomic assemblies [6] and is divergent enough from other synthetic community members to permit unambiguous read mapping, and thus act as a proxy for a clonal experiment.

From the contact map of real HiC data (figure 7a), it can be seen that the rates of intra-chromosomal and inter-chromosomal interactions are roughly equivalent in magnitude. Across the eight chromosomes of *S. stipitis*, there is significant uniformity in the degree of physical intimacy within and between all chromosomes. The subtleties of this chromosomal organisation reveals a self-similar “fuzzy-x” pattern repeated between all chromosomes across the contact map. The convergence point within the pattern is attributed to centromere-SPB binding and has been used to predict centromere locations [38]. It has been shown that the physical constraints generated from the interaction of centromeres to the spindle pole body (SPB) and telomeres to the nuclear envelope are sufficient to explain a number of experimental observations in real data [12, 39]. As Sim3C was derived from study of bacterial datasets, our simulation model does not currently include a notion of these higher organism physical constraints. Consequently, the contact map derived from simulated traditional HiC sequencing elicits a flat field (figure 7b), where the intensity variation that does exist is a byproduct of aforementioned factors such as mappability and cut-site density. For the runtime parameters employed, the rate of intra-chromosomal contact is higher than that of inter-chromosomal, making clear the boundaries between the eight chromosomes (figure 7b). Though our model is presently incomplete for higher organisms, there remains a potential utility as an analytical or simply observational prior.

**Figure 7.**
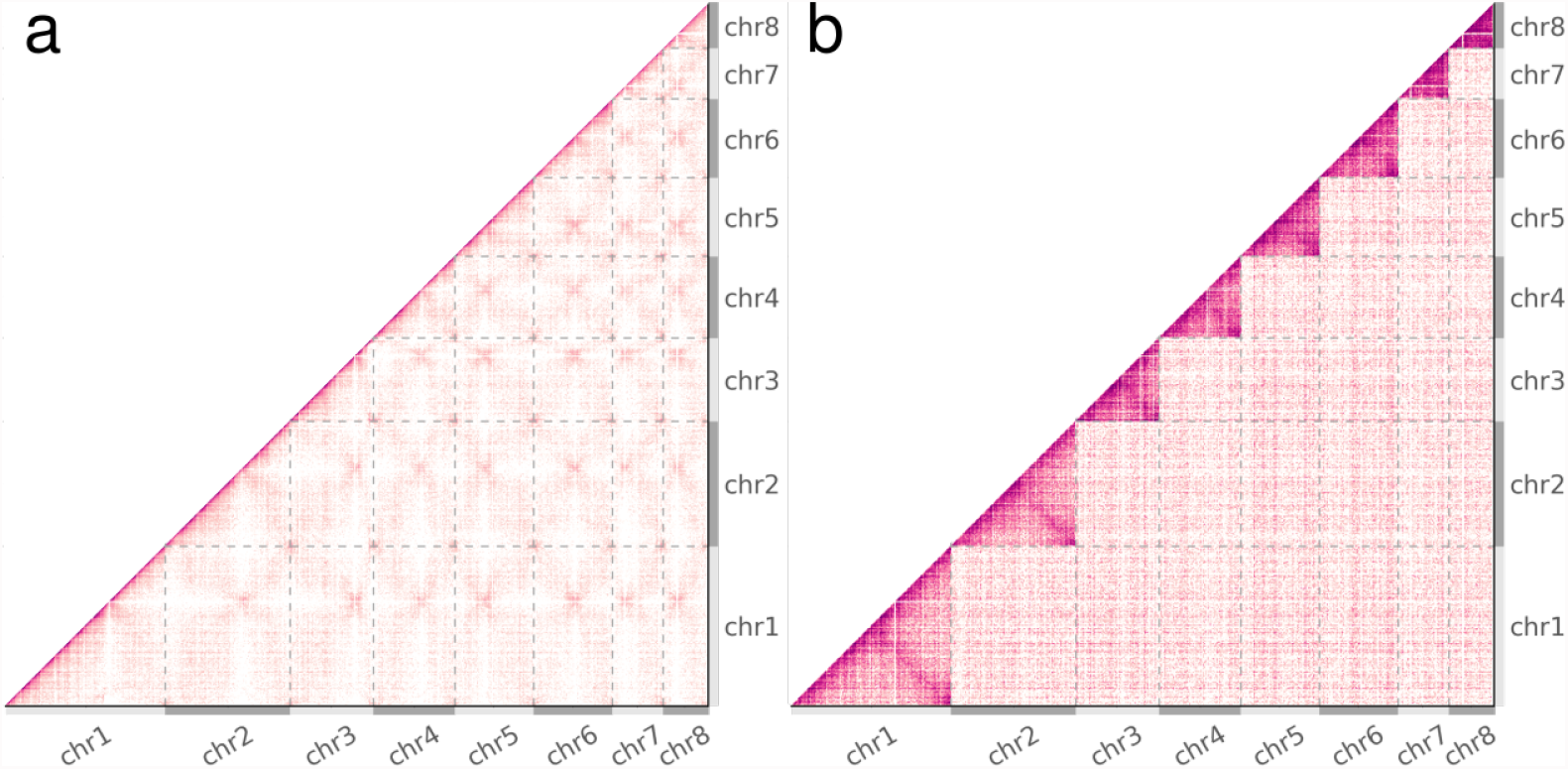
Eukaryotic contact maps. Observed HiC interactions (**a**) real and (**b**) simulated data from the eight chromosome genome of the budding yeast *Scheffersomyces stipitis* CBS 6054 [6]. Grey dashed lines and alternating light and dark grey axes demarcate the boundaries between chromosomes. (**b**) Simulated data elicits a flat field and the clearly evident higher rate of intra- to inter-interactions makes for easily observable chromosomal boundaries within the map. (**a**) Contrastingly for real data, the similar rates of intra-chr and inter-chr interactions reveals the physical constraints imposed by centromere-SPB tethering on all eight chromosomes [38].

### Metagenomic

In the deconvolution of metagenomes, proximity ligation methods hold great potential as new sources of information and have been investigated by the construction and sequencing of synthetic communities [4, 6, 26]. We selected two previously constructed synthetic bacterial communities, one employing traditional HiC and the other Meta3C (table 1). Intended as “proof of concept” experiments, neither community reflects a real environment, but rather were intended to be easily interpreted and include interesting features, such as: range of GC, single and multi- chromosomal genomes and strain-level divergence. The HiC community involved five genotypes from four species, one genome of two chromosomes (*B. thailandensis*), *E. coli* strains BL21 and K12 (Average Nucleotide Identity, ANI 99%) and a wide overall GC range of 37-68% (table 2). Of lower complexity, the Meta3C community involved three genomes from three species, included one genome of two chromosomes (*V. cholerae*) and had a narrower GC range of 44-51% (table 3). Relative to the single genome experiments above, a lower depth of sequencing resulted in a lower overall contact map intensity (figure 8). This is particularly the case for Meta3C, where, by the nature of the method, a large proportion (approx. 99%) of the sequencing yield is in reality conventional WGS read-pair data [26]. As a direct result, in binning the Meta3C dataset, there were insufficient counts to fully establish finer detail within the contact maps, leaving a smoother appearance.

**Figure 8.**
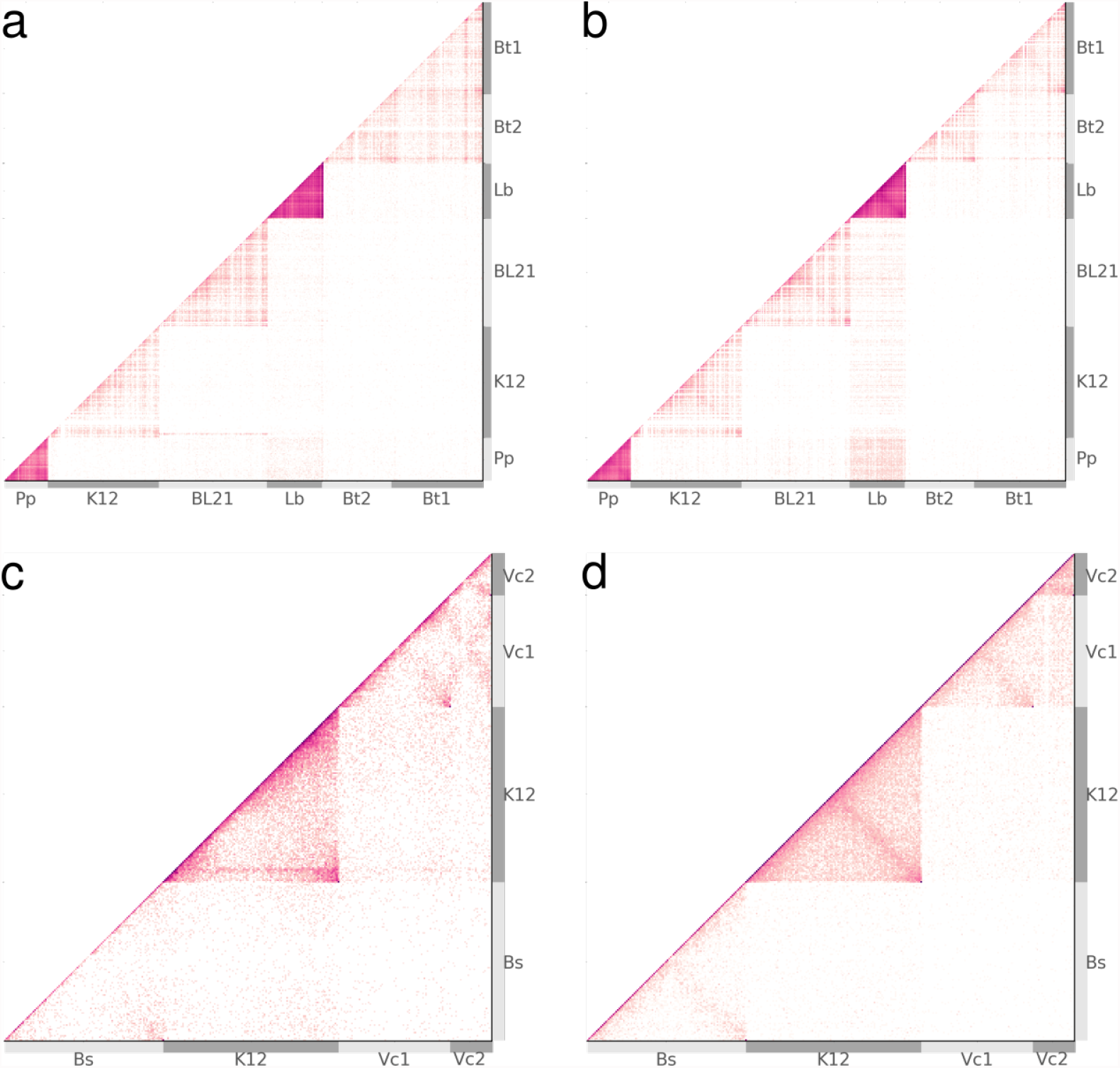
Metagenomic contact maps. Observed HiC interactions (**a**) real and (**b**) simulated data from the eight chromosome genome of *Scheffersomyces stipitis* CBS 6054 [6]. Grey dashed lines and alternating light and dark grey axes demarcate the boundaries between chromosomes. (**b**) Simulated data elicits a flat field and the clearly evident higher rate of intra- to inter- interactions makes for easily observable chromosomal boundaries within the map. (**a**) Contrastingly for real data, the similar rates of intra-chr and inter-chr interactions reveals the physical constraints imposed by centromere-SPB tethering on all eight chromosomes [38].

**Table 2.** Synthetic HiC community. A synthetic community used to demonstrate the utility of HiC sequencing data in resolving a microbial metagenome [4]. It is composed of 5 bacteria, including two closely related strains (*E. coli* K12 and BL21), a genome with two plasmids (*L. brevis)* and a two-chromosome genome (*B. thailandensis*). A is relative abundance, *n_cpy_* is copy number, *n_x_* is number of restriction sites, and *m* = *n_x_/n*_0_ is match quality between chromosome and enzyme choice: *m* < 1 is worse, *m* > 1 is better.

| Name | Replicons | Accession | Chr abbr. | $A$ | $n_{cpy}$ | %GC | $n_x$ | $m$ |
| --- | --- | --- | --- | --- | --- | --- | --- | --- |
| *Burkholderia thailandensis* E264 | 2 | NC_007651<br>NC_007650 | Bt1<br>Bt2 | 0.054 | 1 | 67.29<br>68.07 | 225<br>144 | 0.24<br>0.20 |
| *Escherichia coli* BL21 | 1 | NC_012892 | BL21 | 0.242 | 1 | 50.83 | 508 | 0.46 |
| *Escherichia coli* K12 DH10B | 1 | NC_010473 | K12 | 0.166 | 1 | 50.78 | 568 | 0.50 |
| *Lactobacillus brevis* ATCC 367 | 3 | NC_008497<br>NC_008498<br>NC_008499 | Lb<br>-<br>- | 0.436 | 1 | 46.22<br>38.64<br>38.51 | 629<br>3<br>16 | 1.12<br>0.92<br>1.84 |
| *Pediococcus pentosaceus* ATCC 25745 | 1 | NC_008525 | Pp | 0.102 | 1 | 37.36 | 863 | 1.93 |

**Table 3.** Synthetic Meta3C community. A synthetic community used to demon-strate the utility of Meta3C sequencing data in resolving a microbial metagenome [25, 26]. It is composed of three bacteria with one possessing two chromosomes. *A* is relative abundance, *n_cpy_* is copy number, *n_x_* is number of restriction sites, and *m* = *n_x_/n*_0_ is match quality between chromosome and enzyme choice: *m* < 1 is worse, *m* > 1 is better.

| Name | Replicons | Accession | Chr abbr. | $A$ | $n_{cpy}$ | %GC | $n_x$ | $m$ |
| --- | --- | --- | --- | --- | --- | --- | --- | --- |
| *Bacillus subtilis* subsp. subtilis str. 168 | 1 | NC_000964 | Bs | 0.123 | 1 | 43.51 | 14529 | 0.88 |
| *Escherichia coli* str. K-12 substr. MG1655 | 1 | NC_000913 | K12 | 0.562 | 1 | 50.79 | 24311 | 1.34 |
| *Vibrio cholerae* O1 biovar El Tor str. N16961 | 2 | NC_002505 | Vc1 | 0.332 | 1 | 47.70 | 5909 | 0.51 |
|  |  | NC_002506 | Vc2 |  |  | 46.91 | 1802 | 0.43 |

As with single-genome experiments, metagenomic contact maps are locally modulated by factors such as mappability and cut-site density. Importantly now for metagenomes, the factors of relative abundance and GC content interact to alter the observed intensity of each chromosome within the contact map.

As a first approximation and assuming agreement in nucleotide sampling frequency, we expect *n*_0_ = *L/*4*^λ^* recognition sites for an enzyme of site length *λ* and DNA sequence length *L*. The degree to which an enzyme and DNA sequence deviate from this estimate could be described as how well they match, *m* = *n_x_/n*_0_. Poorer quality matches (*m* < 1) occur when an enzyme's recognition site is underrepresented, while conversely, better quality matches (*m* > 1) describe a situation of more recognition sites than expected.

When multiple chromosomes are taken as a community, the relative proportion of sites from each represents an observational bias when conducting 3C-based experiments. For community *C*, the number of sites *n_x_* from chromosome *x* determines the number of potential PL pairings *N_x_* within *C* which involve *x* (equation 8). The number of intra-chromosomal and inter-chromosomal potential pairs thus respectively vary quadratically and linearly with *n_x_*. Regarding the process of observing a PL event (read-pair) from the community as a random draw with replacement, and the selection pool as comprised of all potential events from all chromosomes, then variation in match quality constitutes a per-chromosome bias. In real laboratory experiments, the composition of the selection pool is further modified by variation in other factors, such as cellular lysis efficiency, unintended DNA fragmentation and relative abundance. In particular, when relative abundances A are introduced, the odds of observing a PL event involving chromosome *x* is then proportional the product *p_x_* ∝ *A_x_ N_x_ / N_C_*. Although the processes of intra-chromosomal, inter-chromosomal, and inter-cellular (spurious) ligation are treated independently in our simulation model, in this manner, per-chromosome intensity (observation rate of chromosome *x*) can vary significantly within a metagenome.

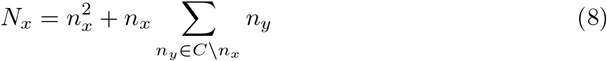

Though the original laboratory experiments reported by Beitel et al. 2014 and Marbouty et al. 2014 intended o create synthetic communities with uniform relative abundances, in practice each possesses a non-uniform profile. The variation in GC content is largest for the HiC experiment and together with non-uniform relative abundances produces a wide range of chromosome intensity for both real and simulated data (figure 8a,b). For both the real and simulated HiC maps, the frequent observation of PL events involving *P. pentosaceus* (Pp) and *L. brevis* (Lb), suggests the possibility that inter-cellular interaction is significant. Within the simulated map at least, inter-cellular pairs are produced exclusively through the process of spurious ligation (noise) and are observed at a higher rate than in the real data, indicating that as expected, spurious ligation rates across species are correlated with their relative abundances.

Further for the HiC data, the two-chromosome genome of *B. thailandensis* (Bt1, Bt2) (figure 8a) has a greater rate of inter-chromosomal interaction than expected from comparing it to simulation (figure 8b). Meanwhile, the clear delineation of *E. coli* strains BL21 and K12 (*ANI* > 99%), with little inter-cellular signal, helps to support the notion that the inter-chromosomal interactions observed between *B. thailandensis* chromosomes (*ANI* ≃ 83%) are real and not a by-product of inadequate filtering.

## Methods

### Reference Data

To compare Sim3C against real experiments, we obtained previously published experimental read-pair datasets (table 1) and their accompanying reference genomes (tables 2, 3) from public archives. In the case of the single genome project of *Caulobacter crescentus* CB15 [18], sequencing data derived from untreated swarmer cells was chosen and the laboratory strain *C. crescentus* NA1000 (acc: NC 011916) was used as the reference genome. For the yeast genome, the completed eight chromosome genome of *Scheffersomyces stipitis* CBS 6054 was used as a reference (acc: PRJNA18881) and the respective reads were extracted from the MY16 yeast synthetic metagenome [6] by direct mapping with BWA MEM. Extraction by mapping in isolation was employed as *S. stipitis* was the second furthest phylogenetically removed yeast in the synthetic community and was the most contiguous (N50: 60kbp) from the whole synthetic community de novo metagenomic WGS assembly.

### Read Generation

Experimental parameters used in read simulation were set to agree as closely as reasonably possible to the respective real experiments, employing the same read length and restriction enzyme (table 1). In each experiment, the published fragment size range was approximated by a normal distribution (table 4). For ease of reproducibility, a single random seed (`1234`) was used in all simulations. As our intent was primarily to demonstrate functionality, rates of inter-chromosomal and spurious events were adjusted per-experiment only through a qualitative process. For simulation of metagenomic datasets, relative abundances were estimated by mapping real experimental reads to the respective reference genomes. From each real experiment, the off-diagonal weight of the resulting contact map was used to calibrate the amount of simulated sequencing required to achieve roughly equivalent intensity (table 4). Both real and simulated read-pair datasets were mapped to their respective reference genomes using BWA MEM (v0.7.15-r1140) [20]

**Table 4.** Runtime simulation. Parameters supplied to Sim3C during read generation.

| Experiment | Insert $\mu$<br>(bp) | Insert $\sigma$<br>(bp) | Anti<br>rate | Spurious<br>rate | Trans<br>rate | Reads<br>($\times 10^6$) |
| --- | --- | --- | --- | --- | --- | --- |
| Beitel et al | 300 | 50 | 0.2 | 0.05 | 0.1 | 7 |
| Burton et al | 400 | 50 | 0.2 | 0.5 | 0.15 | 1.5 |
| Le et al | 400 | 100 | 0.2 | 0.2 | 0.1 | 22 |
| Marbouty et al | 600 | 100 | 0.2 | 0.2 | 0.2 | 7.5 |

### Contact Maps

Contact maps were produced using our own tool (`contact_map.py`), where heatmap intensity was plotted as log-scaled observational frequency. All aligned reads were subject to the same basic filtering criteria: BWA MEM mapq > 5 and alignment length ≥ 50% of read length, with the added restriction that read alignments must have begun with a match. For methods which employed a restriction enzyme (traditional HiC, Meta3C), we constrained the maximum allowable distance from an aligned read to the nearest upstream cut-site. Calculated per chromosome, this distance constraint could not exceed two-fold the median cut-site spacing. Rather than simply delete the primary diagonal for the sake of reducing the displayed dynamic range in figures, we instead to reduced its intensity by categorizing properly paired reads with an estimated fragment size of less than 2 of the reported mean as being conventional WGS (non-PL) reads and ignored them. The resolution of contact maps was adjusted between experiments so as to present a sufficiently bright image without undue loss of resolution. The contact map bin sizes employed were: 10000 bp for the single bacterial genome, 25000 bp for the yeast genome and 40000 bp for the HiC and Meta3C metagenomes (tables 2, 3).

## Availability of supporting source code and requirements

- Project name: sim3C
- Project homepage: https://github.com/cerebis/sim3C
- Operating system: Platform independent
- Programming languages: Python 2.7
- License: GNU GPL v3

## Declarations

### List of abbreviations

- IPC - interprocess communication
- PL - proximity ligation
- WGS - whole genome shotgun
- CID - chromosomal interaction domain
- TAD - topologically associated domain
- *Bern*(*x*) - Bernoulli distribution
- *U* (*x*) - uniform distribution
- *N* (*µ, σ*) - normal distribution
- *cis* - intra-chromosomal
- *trans* - inter-chromosomal

## Ethics approval and consent to participate

Not applicable

## Funding

This work was supported under Australian Research Council's Discovery Projects funding scheme (project number: LP150100912, CI: Djordjevic, SP). The NeCTAR Research Cloud is an Australian Government project conducted as part of the Super Science initiative and financed by the and the Education Investment Fund (EIF) and National Collaborative Research Infrastructure Strategy (NCRIS).

- https://www.education.gov.au/education-investment-fund
- https://www.education.gov.au/national-collaborative-research-infrastructure-strategy-ncris

### Authors contributions

MD designed and implemented Sim3C and wrote the manuscript and prepared figures. AD assisted in the design and contributed to the manuscript.

## Acknowledgements

We thank Steven P. Djordjevic for his support and helpful discussions. This work was supported by the AusGEM initiative, a collaboration between the NSW Department of Primary Industries and the ithree institute. We acknowledge the use of computing resources from the NeCTAR Research Cloud, the QCIF and the UTS eResearch Group.

- http://www.nectar.org.au
- http://www.qcif.edu.au
- https://eresearch.uts.edu.au

